# Regulation of adipocyte dedifferentiation at the skin wound edge

**DOI:** 10.1101/2023.11.22.568302

**Authors:** Longbiao Yao, Sunhye Jeong, Hae Ryong Kwon, Lorin E. Olson

**Author notes:** **Corresponding Author:** Lorin E. Olson, Oklahoma Medical Research Foundation, 825 NE 13^th^ Street, Oklahoma City, OK 73104.

## Abstract

Adipocytes have diverse roles in energy storage and metabolism, inflammation, and tissue repair. Mature adipocytes have been assumed to be terminally differentiated cells. However, recent evidence suggests that adipocytes retain substantial phenotypic plasticity, with potential to dedifferentiate into fibroblast-like cells under physiological and pathological conditions. Here, we develop a two-step lineage tracing approach based on the observation that fibroblasts express platelet-derived growth factor receptor alpha (*Pdgfra*) while adipocytes express Adiponectin (*Adipoq*) but not *Pdgfra*. Our approach specifically traces *Pdgfra*^+^ cells that originate from *Adipoq*^+^ adipocytes. We find many traced adipocytes and fibroblast-like cells surrounding skin wounds, but only a few traced cells localize to the wound center. In agreement with adipocyte plasticity, traced adipocytes incorporate EdU, downregulate Plin1 and PPARγ, and upregulate αSMA. We also investigate the role of potential dedifferentiation signals using constitutively active PDGFRα mutation, *Pdgfra* knockout, or *Tgfbr2* knockout models. We find that PDGF and TGFβ signaling both promote dedifferentiation, and PDGFRα does so independently of TGFβR2. These results demonstrate an intersectional genetic approach to trace the hybrid cell phenotype of *Pdgfra*^*+*^ adipocytes, which may be important for wound repair, regeneration and fibrosis.

## INTRODUCTION

Mammalian white adipocytes are lipid storing cells found throughout the body. In many locations they occur as discrete subcutaneous and visceral depots, but they also reside along blood and lymphatic vessels, within and around muscle, and between organs. White adipocytes undergo dramatic changes in volume to accommodate lipid storage needs (Sakers et al., 2022). Lipolysis is the process by which adipocytes break down their stored triglyceride and release fatty acid, resulting in adipocyte shrinkage. When there is a positive energy balance, adipocytes take up fatty acid and convert it into triglyceride to expand their lipid droplet. Adipocytes develop from adipocyte progenitors (AP), which are fibroblast-like cells in development and adulthood. Cell sorting and lineage tracing approaches have identified APs as positive for platelet-derived growth factor receptor-α (PDGFRα) or - β (PDGFRβ), along with other markers (Berry and Rodeheffer, 2013; Rodeheffer et al., 2008; Tang et al., 2008). When isolated from mice and cultured in vitro, these cells form colonies and differentiate into adipocytes. When transplanted into fatless mice, they populate fat pads with functional adipocytes (Rodeheffer et al., 2008). Dermal fat contains its own resident population of functional PDGFRα^+^ APs (Driskell et al., 2013; Rivera-Gonzalez et al., 2016; Zhang et al., 2016). Single cell RNA sequencing and ATAC sequencing from newborn mouse skin supports the conclusion that APs and adipocytes are a subset of the fibroblast population (Thompson et al., 2022).

Dermal fat in mice is separate and distinct from subcutaneous adipose tissue (Wojciechowicz et al., 2013; Zwick et al., 2018). Dermal adipocytes store lipid, but they also have anti-microbial properties and an intimate relationship with hair growth (Festa et al., 2011; Zhang et al., 2016; Zhang et al., 2015). Recently, dermal adipocytes were shown to participate in skin wound repair by activating lipolysis and reverting into fibroblast-like cells (Schmidt and Horsley, 2013; Shook et al., 2020). This process, called dedifferentiation or adipocyte myofibroblast transition (AMT), is distinct from mere adipocyte shrinkage as occurs during lipolysis (Song and Kuang, 2019; Zhu et al., 2022). Dedifferentiation involves loss of adipocyte molecular markers *Adipoq* (adiponectin), *Plin1* (perilipin 1), and *Pparg* (PPARγ), re-expression of AP/fibroblast markers *Pdgfra* and *Pdgfrb*, and re-acquisition of the ability to proliferate. Some adipocyte-derived fibroblasts migrate into the wound bed and join the population of fibroblasts and myofibroblasts that create the scar tissue (Shook et al., 2020). Lipid release attracts macrophages to the wound and fuels angiogenesis in the wound bed. Some adipocyte-derived cells that localize to the scar may become myofibroblasts. Many cytokine and growth factor receptors are upregulated by dedifferentiated adipocytes in skin wounds, including receptors for PDGF and TGFβ (Shook et al., 2020), but whether these signals regulate dedifferentiation is unknown. More recently, the process of adipocyte dedifferentiation has been questioned by some investigators (Kalgudde Gopal et al., 2023).

In this study, we wanted to improve the labeling and visualization of adipocyte dedifferentiation. If large numbers of adipocytes undergo dedifferentiation and lineage expansion, this would be highly significant for our understanding tissue homeostasis, regeneration, and fibrosis. Current lineage tracing approaches using Adipoq-CreER or Adipoq-rtTA/TRE-Cre (AdipoChaser) are not specific to dedifferentiated adipocytes because they label mature adipocytes with high efficiency (Jeffery et al., 2014; Wang et al., 2013b). As a result of all adipocytes being strongly labeled, it is challenging to identify the small number of cells that change gene expression and morphology. Therefore, to restrict labeling to the subset of adipocytes specified for dedifferentiation, as defined by re-expression of the *Pdgfra*, we combined *Adipoq-CreER* with *Pdgfra-loxP-Stop-loxP-Flp*^*o*^ (*Pdgfra*^*Flp*^) and *R26-FRT-Stop-FRT-tdTomato* mice. These three components create a tamoxifen-regulated genetic tracing system where Tomato is expressed specifically in *Adipoq*^*+*^ adipocytes if they also express *Pdgfra*. The strength of this system is that it does not label adipocytes at homeostasis, but selectively traces those that express *Pdgfra*. We use this system to investigate dedifferentiation of dermal adipocytes during wound healing.

## RESULTS

### Specific labeling of Pdgfra^+^ adipocytes at the wound edge and center

In adult mice, *Adipoq* is expressed by mature adipocytes but not preadipocytes or adipocyte progenitors (APs), and conversely, *Pdgfra* is expressed by fibroblasts and APs but not mature adipocytes (Berry and Rodeheffer, 2013; Emont et al., 2022; Lee et al., 2012; Roh et al., 2017; Wang et al., 2010). Previously we characterized *Pdgfra*^*Flp*^ mice with a cassette containing a floxed cDNA encoding *Pdgfra* followed by a *Flp*^*o*^ sequence, which was inserted to replace the endogenous *Pdgfra* gene. Cre recombination in these mice leads to knockout of the *Pdgfra* cDNA and expression of Flp° controlled by the endogenous *Pdgfra* gene (Sun et al., 2020; Yao et al., 2022). These alleles can be combined with any Cre driver and a Flp/FRT-dependent reporter to perform intersectional genetic tracing. Here we combined *AdipoqCreER* with *Pdgfra*^*Flp*^ and *R26*^*fSf-tdTomato*^ to create a tamoxifen-regulated genetic tracing system where Tomato is expressed specifically in *Adipoq*^*+*^ adipocytes that co-express *Pdgfra*. We call this mouse <u>A</u>dipoq-<u>P</u>dgfr<u>α</u>-<u>T</u>racer or APαT (**Figure 1A**).

**Figure 1.**
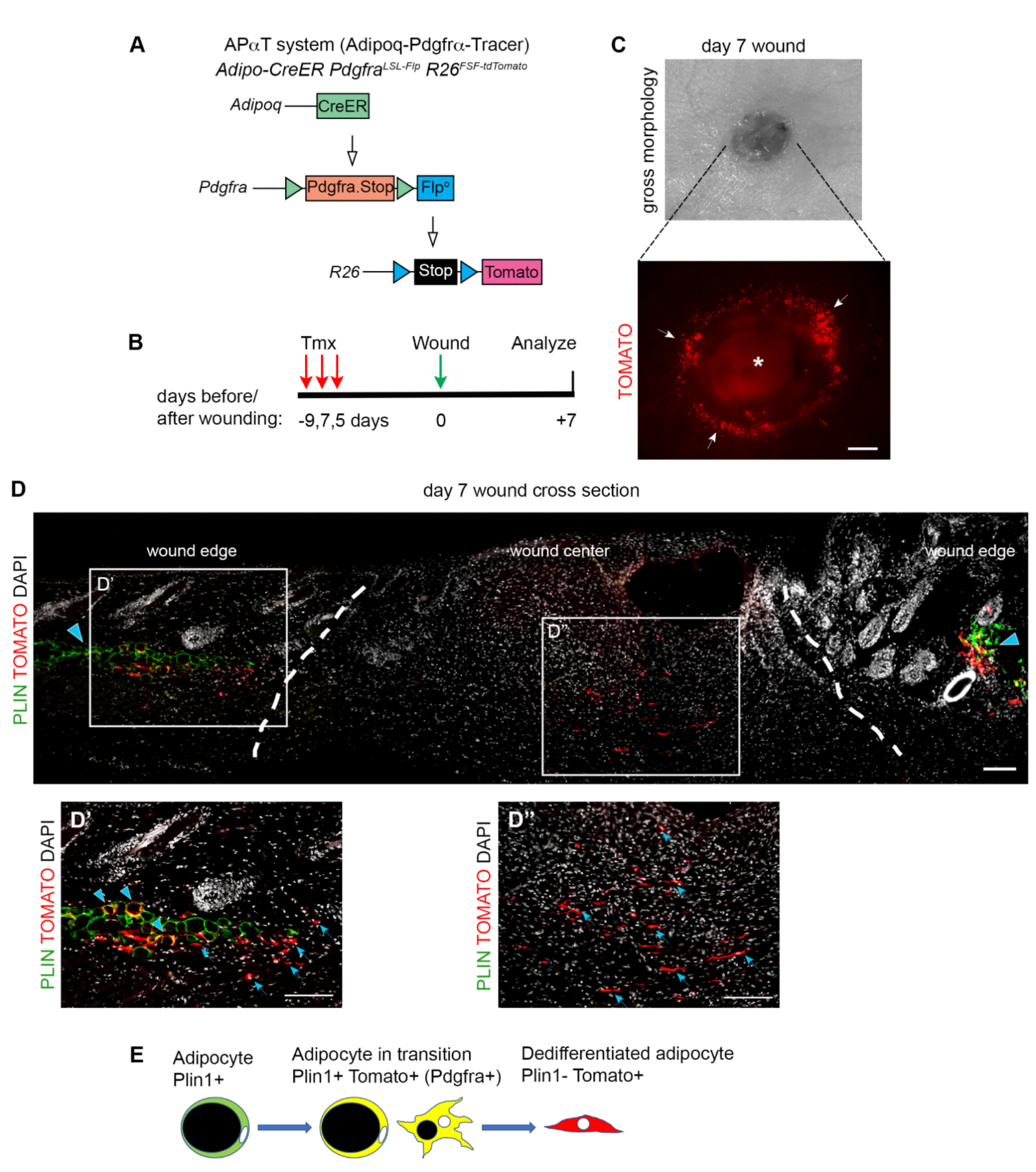
Specific labeling of adipocytes at the wound edge. (A) Schematic of the genetic strategy to label adipocytes that express *Pdgfra*. (B) Timeline for wound experiments. (C) Photograph of a 5mm skin wound on day 7 of healing (top) and a whole mount fluorescent image of a day 7 wound showing Tomato reporter activity specifically in cells at the wound edge (bottom, arrows). Dim autofluorescence is seen at the wound center (asterisk). Scale bar 500 μm. (D) Cross section through a day 7 wound. Plin1^+^ cells are seen at the wound edges (arrowheads) with interspersed Tomato^+^ cells emerging close to the wound edge (dotted line). (D’) Zoom-in highlighting Plin1^+^Tomato^+^ adipocytes (arrowheads) and emerging Tomato^+^ Plin1^neg^ cells (arrows). (D’’) Zoom-in highlighting Tomato^+^ Plin1^neg^ cells at the wound center (arrows). Scale bar 100 μm. (E) Model of the adipocyte-to-fibroblast transition.

We administered tamoxifen to 6-weeks-old APαT mice and then created 5mm excisional wounds in the dorsal skin (**Figure 1B**). We extensively examined unwounded skin and other fat depots for Tomato expression, which would indicate *AdipoqCreER* activity overlapping with *Pdgfra*^*Flp*^ expression at homeostasis. However, we did not detect any Tomato labeling in unwounded dermal adipose tissue or in the major subcutaneous, perigonadal, or interscapular depots (**Figure S1**). Therefore, with a 16-day window of analysis in adult mice at homeostasis, there is no *Pdgfra* expression in *Adipoq*^*+*^ adipocytes, and no conversion of adipocytes into fibroblasts. On the other hand, after 7 days of healing, Tomato^+^ cells formed a ring around the perimeter of the wound when visualized by whole mount and viewed from the underside of the skin (**Figure 1C** arrows). The wound center was usually obscured by autofluorescence (**Figure 1C** asterisk). Therefore, cross-sections of the wound were stained for Plin1 (perilipin 1, an adipocyte marker), which revealed a major population of Tomato^+^ cells at the wound edge near the intact layer of dermal adipocytes. In this area, Plin1^neg^Tomato^+^ cells resembled fibroblasts, while Plin1^+^Tomato^+^ cells had an adipocyte morphology (**Figure 1D**,**D’**). There was also a minor population of Tomato^+^ cells near the wound center. These were Plin1^neg^Tomato^+^ fibroblast-like cells that had moved into the wound bed (**Figure 1D’’**). These data show that APαT does not label all adipocytes, as occurs when *AdipoCreER* is used alone, but instead it specifically labels wound-associated adipocytes and their fibroblast-like derivatives. We postulated a multistep dedifferentiation process where Plin1^+^ adipocytes acquire *Pdgfra* expression, become Tomato^+^, change morphology, and lose Plin1 (**Figure 1E**).

### Characterization of the adipocyte-to-fibroblast transition

Large fields of Tomato^+^ cells were observed at the wound edge and sparse Tomato^+^ cells were observed in the wound center when flat-sections were cut parallel to the plane of the skin instead of typical cross-sections (**Figure 2A - C**). A small number of Tomato^+^ adipocyte-derived cells at the wound center is in agreement with previous reports (Schmidt and Horsley, 2013; Shook et al., 2020), but the large number of Tomato^+^ cells at the wound edge was somewhat unexpected. Shook et al. showed that wound edge adipocytes undergo significant changes in size during the first 1.5 days after healing, and adipocyte lineage cells that migrate into the wound bed upregulate *Pdgfra* (Shook et al., 2020). But our result indicates that *Pdgfra* is upregulated in many adipocytes that never move into the wound bed.

**Figure 2.**
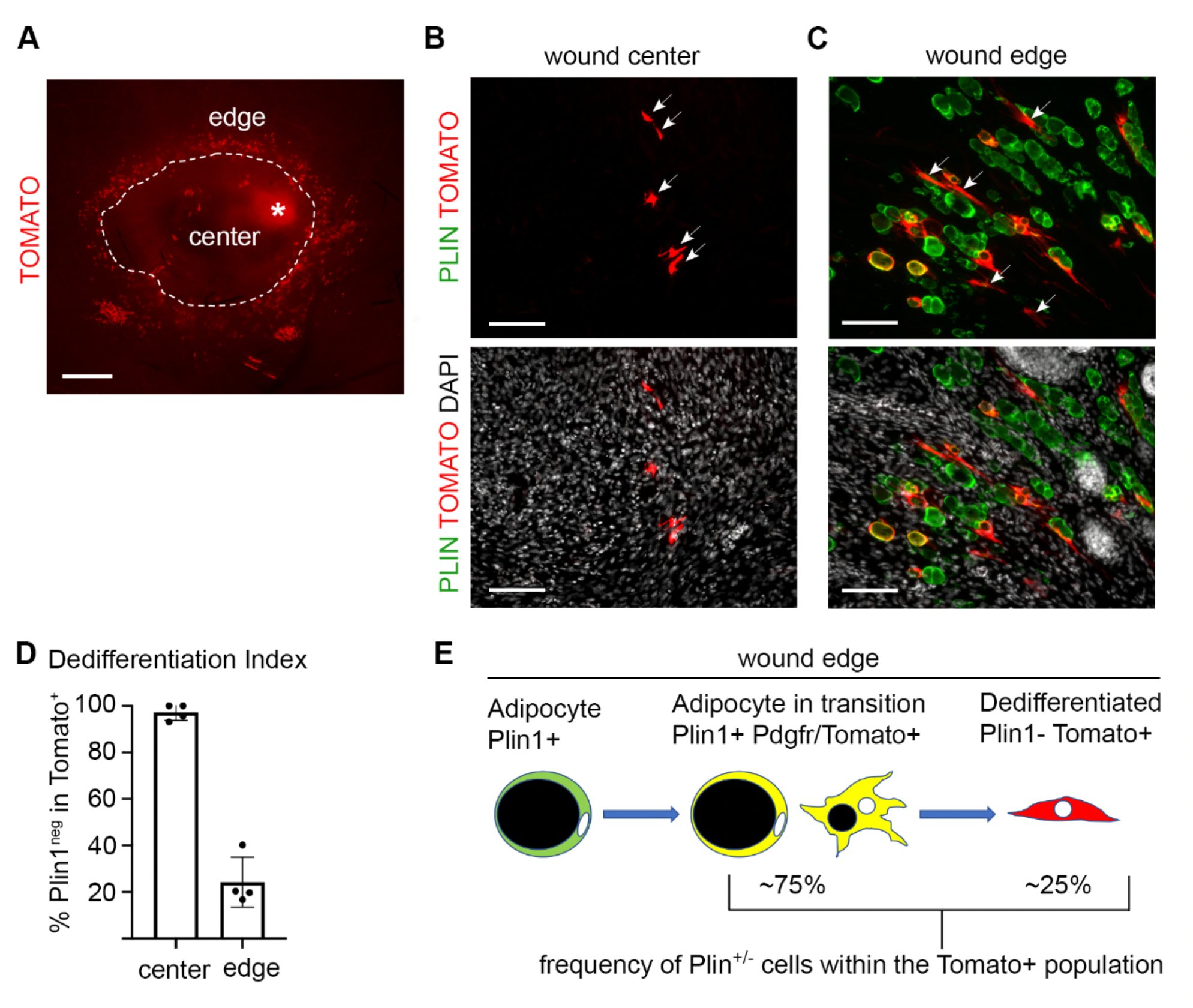
The APαT system traces cells as they cross the adipocyte-to-fibroblast transition. (A) Whole mount fluorescent image of a day 7 wound showing Tomato^+^ in cells at the wound edge. Asterisk, autofluorescence. Scale bar 500 μm. (B, C) Flat sections with Plin1/Tomato co-labeling at the wound center or edge. Arrows indicate Tomato^+^ Plin1^neg^ fibroblast-like cells. Scale bar 100 μm. (D) Quantification of the adipocyte dedifferentiation index, or the percentage of Tomato^+^ cells that are Plin1^neg^. Data are plotted as mean +/- SEM. Each point represents one mouse (n = 4). (E) Summary of the proportion of adipocytes in transition vs dedifferentiated adipocytes.

Tomato^+^ cells at the wound center were always Plin1^neg^ with fibroblast-like morphology (**Figure 2B)**. But the wound edge harbored a mixed population of Plin1^+^Tomato^+^ (yellow) cells with adipocyte or hybrid morphology and Plin1^neg^Tomato^+^ (red) cells with fibroblast-like morphology (**Figure 2C**). To determine the proportions of the two cell types, we calculated a “dedifferentiation index”, which is the number of Plin1^neg^Tomato^+^ fibroblasts divided by the total number of Tomato^+^ cells sampled. The index at the wound center was almost 100% dedifferentiated (Plin1^neg^Tomato^+^) but the wound edge was 24.23 +/- 10.74% dedifferentiated (**Figure 2D**). We conclude that most adipocytes that acquire *Pdgfra* expression remain Plin1^+^ with an adipocyte-like morphology and do not undergo dedifferentiation (**Figure 2E**).

Pparγ is the master transcription factor for adipocytes. It is sufficient to convert fibroblasts into adipocytes and is required for the differentiation and maintenance of adipocytes (Tontonoz et al., 1994; Wang et al., 2013a; Wang et al., 2015). Therefore, Tomato^+^ cells with Pparγ are adipocytes and Pparγ^neg^ cells are no longer adipocytes. Immunofluorescence on APαT wounds showed absence of Pparγ in Tomato^+^ fibroblasts at the wound center, but Pparγ expression was retained in Tomato^+^ cells with adipocyte morphology at the wound edge (**Figure 3A, B**). Loss of Pparγ is further evidence of adipocyte dedifferentiation.

**Figure 3.**
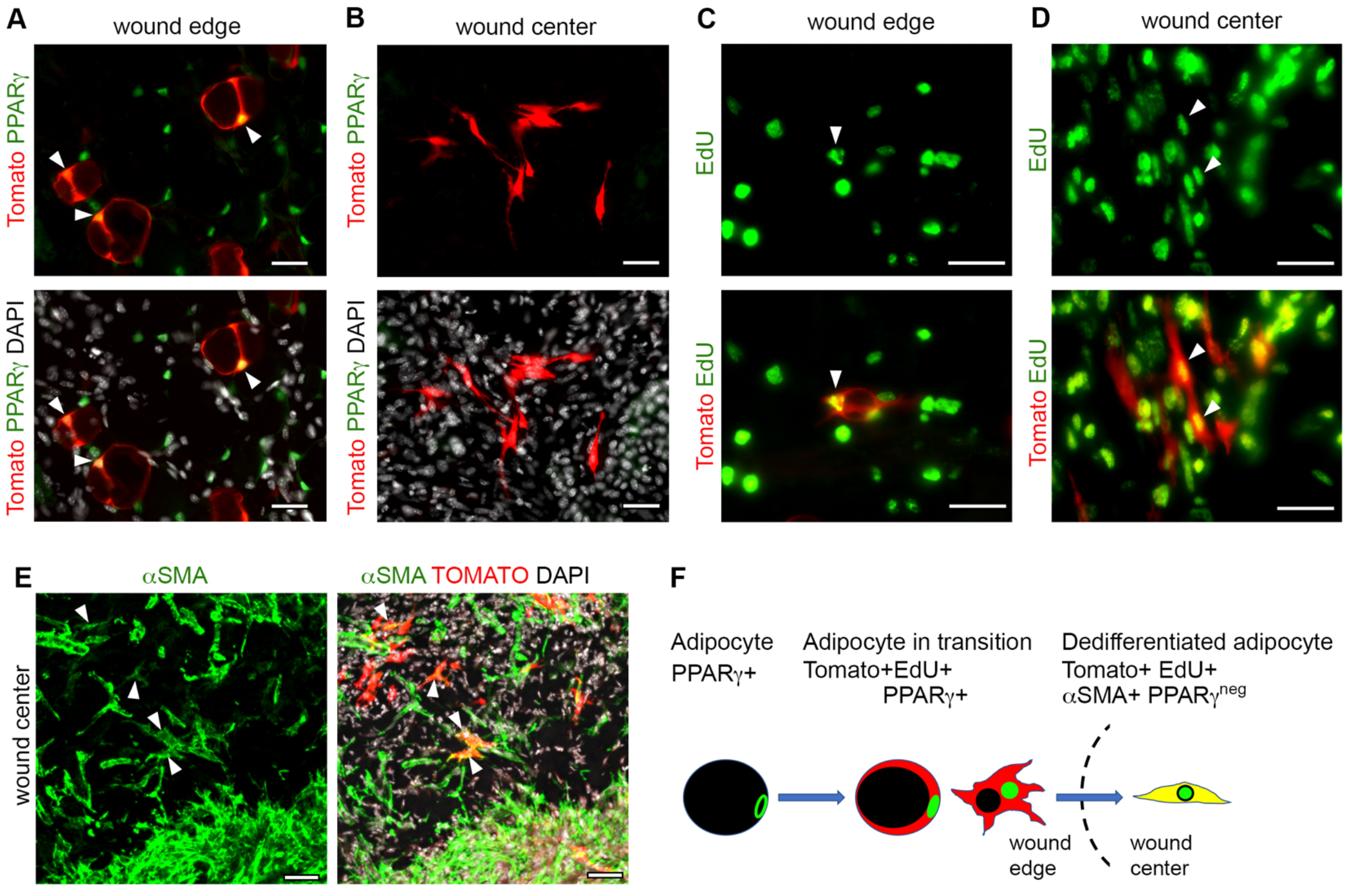
Wound adipocytes replicate DNA, lose PPARγ expression, and acquire αSMA expression. (A, B) Fluorescent images of day 5 wounds showing Tomato^+^ adipocytes at the wound edge with PPARγ (arrowheads in A) or Tomato^+^ dedifferentiated adipocytes at the wound center without PPARγ. (C, D) Fluorescent images of day 5 wounds showing EdU labeling of a Tomato^+^ adipocyte at the wound edge (arrowhead in C) or Tomato^+^ fibroblast-like cells at the wound center (arrowheads in D). (E) Confocal image of the wound center on day 5, showing Tomato^+^ fibroblast-like cells with αSMA. (F) Summary of events across the adipocyte-to-fibroblast/myofibroblast transition. Scale bar 50 μm.

Mature adipocytes cannot divide, so new adipocytes arise from the proliferation and differentiation of APs. However, dedifferentiated adipocytes were reported to incorporate the nucleotide analog EdU, indicating DNA synthesis (Schmidt and Horsley, 2013; Shook et al., 2020). To examine DNA synthesis, we injected APαT mice with EdU on days 4 and 5 of wound healing and harvested tissue 4 hours after the last injection. We saw Tomato^+^EdU^+^ cells with a fibroblast morphology at the wound center as expected (**Figure 3C**), but there were also examples of Tomato^+^EdU^+^ cells at the wound edge with adipocyte morphology (**Figure 3D**). This indicates that DNA replication can occur in adipocytes and is not entirely restricted to preadipocytes or fully dedifferentiated adipocytes.

Dedifferentiated adipocytes that migrate into the granulation tissue may further differentiate into αSMA^+^ myofibroblasts. We stained day 5 wounds for αSMA and observed αSMA^+^Tomato^+^ cells in the wound center (**Figure 3E**). Therefore, a minor portion of the Tomato^+^ fibroblasts in APαT mice reach the wound center and express αSMA, which signifies activation of transcriptional programs governing contractile function. But the contribution of dedifferentiated adipocytes to the overall wound myofibroblasts is very small compared to the total population of myofibroblasts. Together, these results characterize the adipocyte-to-fibroblast transition as a process involving a sequence of events: 1) gain of fibroblast marker PDGFRα and re-entry into the cell cycle, 2) loss of lipid droplet concomitant with loss of adipocyte markers Plin1 and PPARγ, which signifies dedifferentiation, and 3) a fraction of the dedifferentiated cells move to the wound center and express αSMA (**Figure 3F**).

### Regulation of the adipocyte-to-fibroblast transition by PDGFRα and TGFβR2

Having established an assay to label dedifferentiating adipocytes in wound healing, we used APαT mice to investigate signals regulating this process, which are unknown. Upregulation of *Pdgfra* during dedifferentiation suggests that PDGF signaling might promote dedifferentiation. To investigate the functional role of PDGFRα, we generated double APαT mice with two alleles of *Pdgfra*^*Flp*^ to enable inducible *Pdgfra* knockout (*Pdgfra*-iKO) (**Figure 4A**). We also generated constitutively active APαT mice (*Pdgfra*-iGOF) using the *Pdgfra*^*K*.*Flp*^ allele that expresses a gain-of-function mutant form of PDGFRα with Flp° (Sun et al., 2020; Yao et al., 2022) (**Figure 4B**). After tamoxifen treatment and wounding, Tomato^+^ cells appeared at the wound edge of both genotypes. Wounds on *Pdgfra*-iGOF mice displayed a brighter ring with more Tomato^+^ cells (**Figure 4D**) than either *Pdgfra*-iKO (**Figure 4C**) or control APαT wounds (**Figure 2A**), which is consistent with cell proliferation typically induced by PDGFRα gain of function (Olson and Soriano, 2009). We quantified Plin1 staining of Tomato^+^ cells to generate a dedifferentiation index. In *Pdgfra*-iKO wounds, dedifferentiation occurred in ∼10% of Tomato^+^ cells at the wound edge, significantly lower than APαT wounds. Dedifferentiation was strikingly high in *Pdgfra*-iGOF wounds, with ∼90% of Tomato^+^ cells registering as Plin1^neg^Tomato^+^ with a fibroblastic morphology (**Figure 4E - G**). These results show that PDGFRα is needed for dedifferentiation and active signaling promotes dedifferentiation (**Figure 4H**).

**Figure 4.**
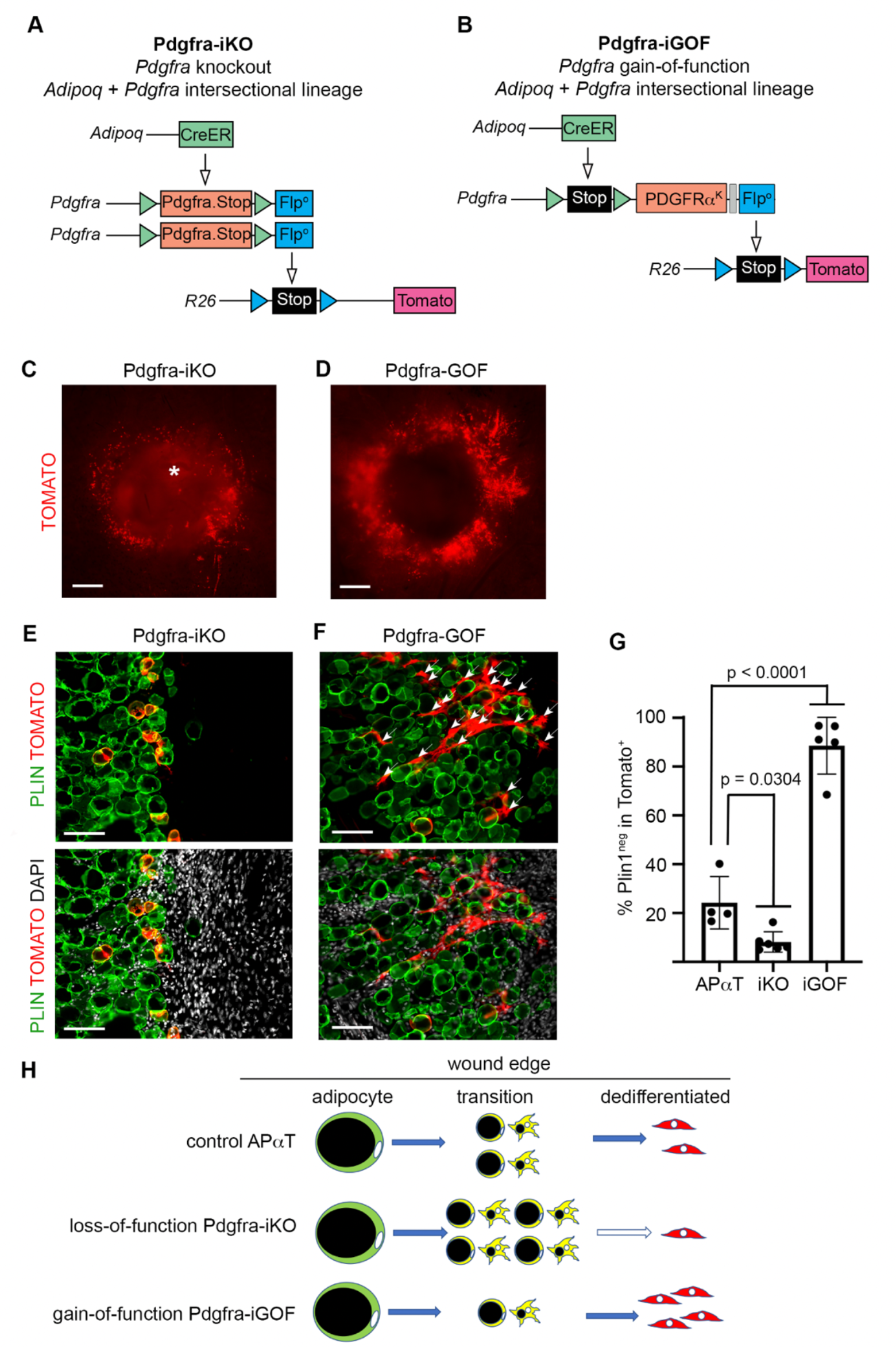
PDGFRα signaling is necessary for and promotes adipocyte dedifferentiation. (A, B) Schematic of the genetic strategy to label dedifferentiating adipocytes combined with *Pdgfra* deletion (Pdgfra-iKO) or with *Pdgfra* gain-of-function (Pdgfra-iGOF). (C, D) Whole mount of a day 7 wound showing Tomato^+^ in cells at the wound edge, which are more abundant in Pdgfra-iGOF. Asterisk, autofluorescence. Scale bar 500 μm. (E, F) Plin1/Tomato co-labeling at the wound edge with Pdgfra-iKO or Pdgfra-iGOF. Arrows indicate Tomato^+^ Plin1^neg^ fibroblast-like cells (arrows in F). Scale bar 100 μm. (G) Quantification of the adipocyte dedifferentiation index for APαT, Pdgfra-iKO and Pdgfra-iGOF. Data are plotted as mean +/- SEM with statistical analysis by one-way ANOVA. Each point represents one mouse (n = 6 iKO, 5 iGOF). Note: the APαT data are from the same mice in figures 2D and 4G. (D) Summary of outcomes from PDGFRα loss-of-function and gain-of-function experiments.

TGFβ is a profibrotic cytokine that suppresses adipocyte identity and promotes fibrotic responses (Roh et al., 2020; Shen et al., 2022). It has been shown to cross-talk with PDGFRα during tissue repair (Contreras et al., 2019). TGFβ receptors are upregulated during the adipocyte-to-fibroblast transition (Shook et al., 2020). To investigate the functional role of the catalytic TGFβ receptor, TGFβR2, we generated APαT mice with inducible deletion of two *Tgfbr2*^*flox*^ alleles (*Tgfbr2*-iKO) or control mice with one *Tgfbr2*^*flox*^ allele (**Figure 5A - B**). We wondered if any Tomato labeling would occur in *Tgfbr2*-iKO wounds, because TGFβ signaling might regulate *Pdgfra* in wound-adjacent adipocytes. However, labeled cells were abundant around *Tgfbr2*-iKO wounds (**Figure 5C - D**). This indicates that *Pdgfra* expression and activation of the APαT system in adipocytes does not require *Tgfbr2*. To determine whether TGFβR2 is functionally involved in the transition from adipocyte to fibroblast, we calculated dedifferentiation at the wound edge. The dedifferentiation index of *Tgfbr2*-iKO wounds was significantly reduced compared to control wounds (**Figure 5E - F**), suggesting that TGFβ signaling is required for dedifferentiation.

**Figure 5.**
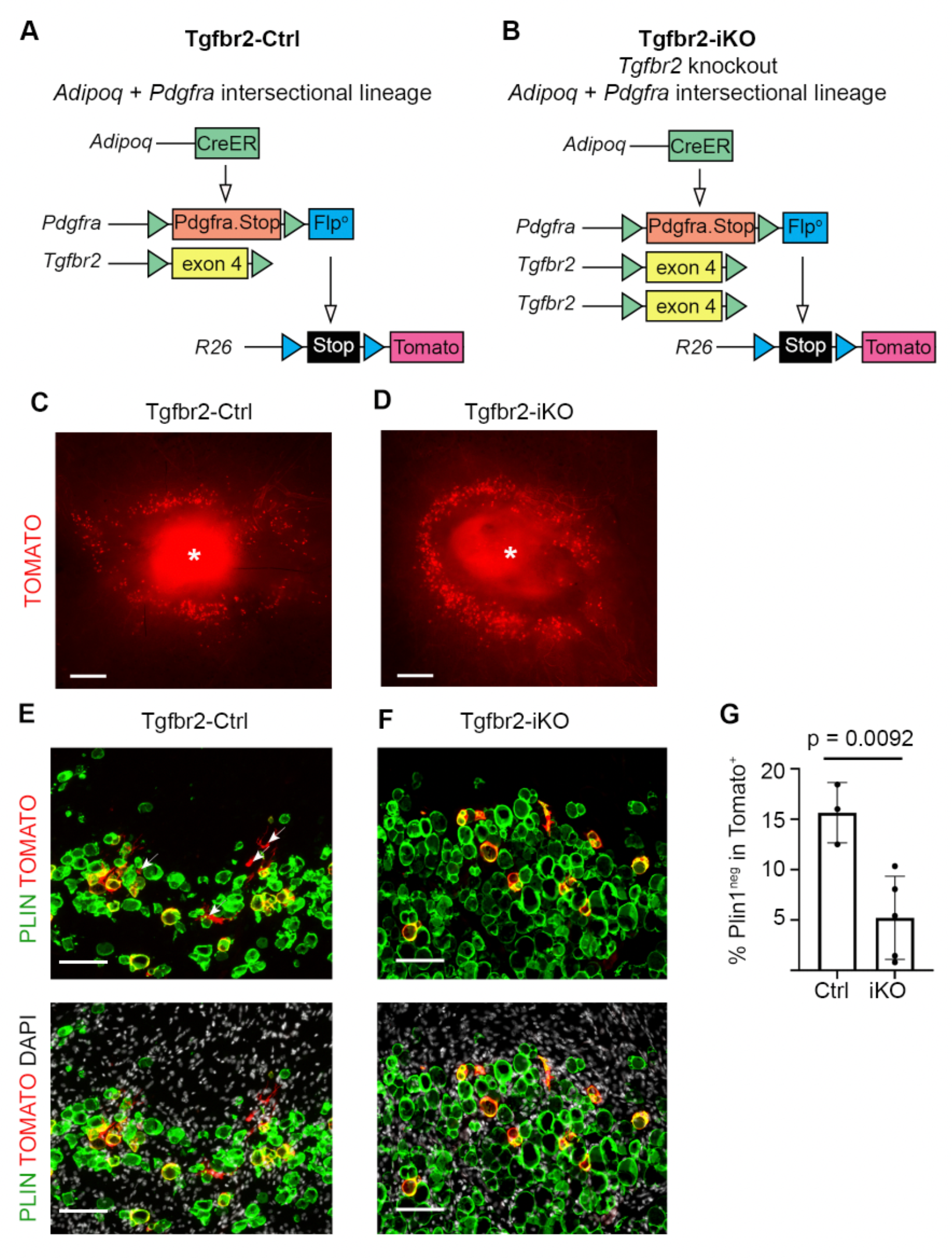
TGFβR2 signaling is necessary for adipocyte dedifferentiation. (A, B) Schematic of the genetic strategy to label dedifferentiating adipocytes combined with *Tgfbr2* deletion (Tgfbr2-iKO). (C, D) Whole mount of a day 7 wound showing Tomato^+^ in cells at the wound edge in both genotypes. Asterisk, autofluorescence. Scale bar 500 μm. (E, F) Plin1/Tomato co-labeling at the wound edge with Tgfbr2-Ctrl or Tgfbr2-iKO. Arrows indicate Tomato^+^ Plin1^neg^ fibroblast-like cells (arrows in E). Scale bar 100 μm. (C) Quantification of the adipocyte dedifferentiation index for Tgfbr2-Ctrl and Tgfbr2-iKO. Data are plotted as mean +/- SEM with statistical analysis by Student’s t-test. Each point represents one mouse (n = 3 Ctrl, 5 iKO).

Because constitutively active *Pdgfra*-iGOF promoted and *Tgfbr2*-iKO inhibited dedifferentiation, we wondered which would dominate in mice with both types of mutations. Hypothetically, the dedifferentiation-promoting effect of constitutive PDGFRα signaling might require TGFβR2 or it might be a TGFβ-independent process. To test this, we generated mice with one or two *Tgfbr2*^*flox*^ alleles plus *Pdgfra*-iGOF and performed tamoxifen treatment and wound procedures as before (**Figure 6A – B**). *Pdgfra*-iGOF wounds with *Tgfbr2*-Ctrl or *Tgfbr2*-iKO exhibited abundant Tomato^+^ cells around the wound edge (**Figure 6C – D**). *Pdgfra*-iGOF + *Tgfbr2*-iKO wounds exhibited a high dedifferentiation index (∼85%) that was close to *Pdgfra*-iGOF + *Tgfbr2*-Ctrl (**Figure 6E - G**) and very different from *Tgfbr2*-iKO (**Figure 5G**). This indicates that the ability of PDGFRα signaling to promote dedifferentiation does not require TGFβR2 (**Figure 6H**).

**Figure 6.**
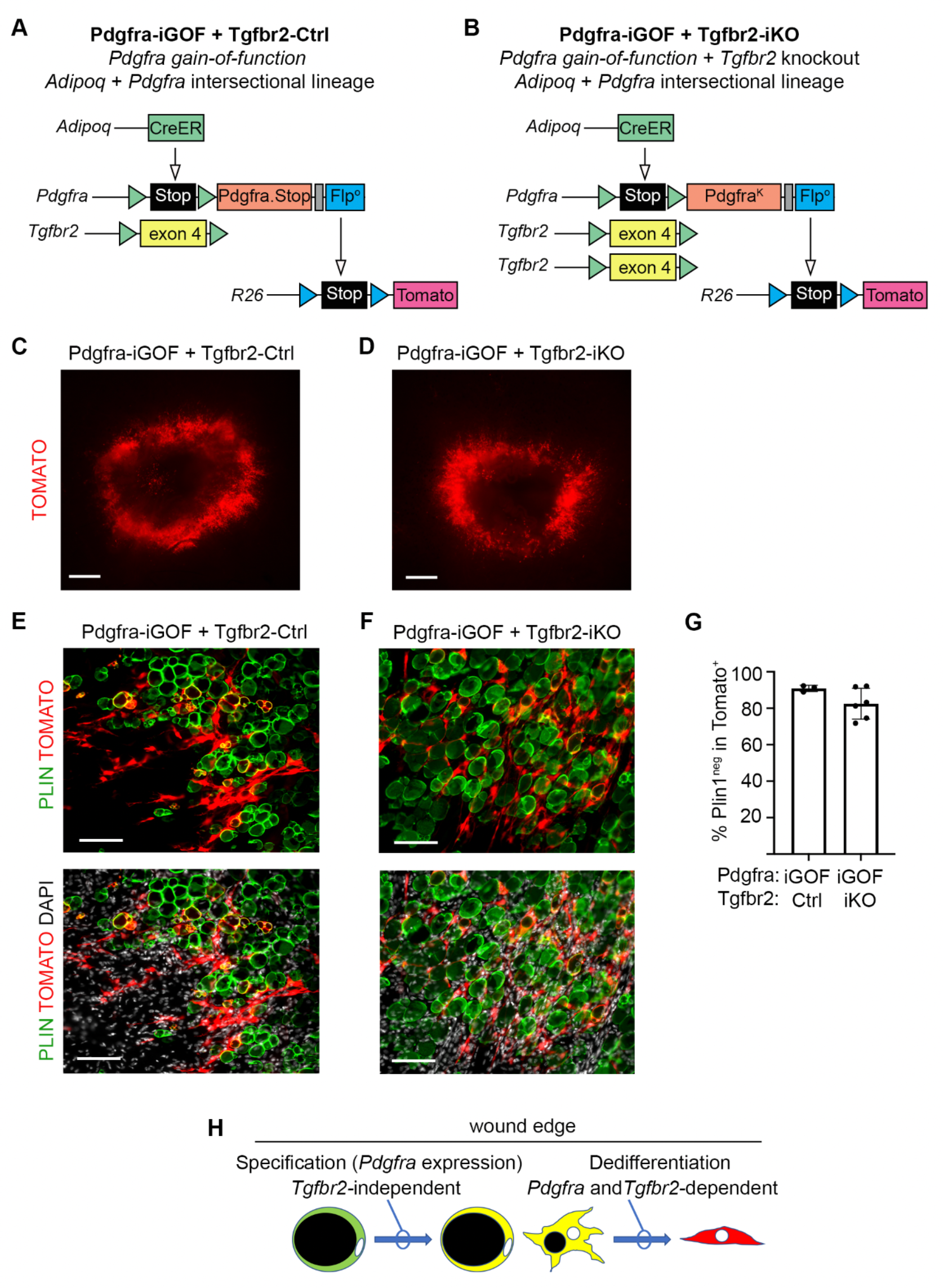
PDGFRα-induced dedifferentiation does not require TGFβR2. (A, B) Schematic of the genetic strategy to label PDGFRα-induced dedifferentiating adipocytes with or without *Tgfbr2* deletion. (C, D) Whole mount of a day 7 wound showing Tomato^+^ cells at the wound edge in both genotypes. Scale bar 500 μm. (E, F) Plin1/Tomato co-labeling at the wound edge with Pdgfra-iGOF + Tgfbr2-Ctrl or Pdgfra-iGOF + Tgfbr2-iKO. Scale bar 100 μm. (G) Quantification of the adipocyte dedifferentiation index for Pdgfra-iGOF + Tgfbr2-Ctrl or Pdgfra-iGOF + Tgfbr2-iKO. Data are plotted as mean +/- SEM with statistical analysis by Student’s t-test. Each point represents one mouse (n = 3 iGOF/Ctrl, 6 iGOF/iKO). (H) Summary of the roles of PDGFRα and TGFβR2 in the adipocyte-to-fibroblast transition.

## DISCUSSION

Adipocyte dedifferentiation has received limited attention compared to adipogenesis. But recent studies propose that dedifferentiation occurs in a variety of physiological and pathological conditions, including wound healing, hair cycling, mammary gland remodeling, and tumor invasion (Shook et al., 2020; Wang et al., 2018; Zhang et al., 2019; Zhu et al., 2022). Previous studies that relied on *Adipoq*-driven Cre alone were able to identify and isolate fibroblast-like cells emerging from an adipocyte lineage, but the strong labeling of all adipocytes was a limitation. In this study, we developed genetic tools to trace the subset of mature adipocytes that acquire progenitor marker *Pdgfra*. Under homeostatic conditions, adipocytes do not express *Pdgfra* and cannot proliferate. However, we find that skin wounding potently induces *Pdgfra* in *Adipoq*^*+*^ adipocytes at the wound edge. Consistent with previous studies, we find that dedifferentiation involves DNA replication, but we also provide evidence to suggest that DNA replication occurs in adipocytes before they lose their lipid droplet, which had not been appreciated. These findings identify a new hybrid cell phenotype where PPARγ^+^ lipid-laden adipocytes express fibroblast markers. In APαT mice with no additional genetic manipulations, 25% of the *Pdgfra*^+^ adipocyte population undergoes morphological change and loses Plin1. We consider these cells to be dedifferentiated adipocytes. The remaining 75% of *Pdgfra*^+^ adipocytes that retain Plin1 and adipocyte morphology may be considered as adipocytes that were specified but not committed to dedifferentiation. Our data show that dedifferentiated adipocytes are a minor population within a major population of *Pdgfra*^+^ adipocytes that do not transition into fibroblast-like cells. It was not possible to visualize this majority population in previous studies using *Adipoq*-driven Cre alone.

Understanding the hybrid cell phenotype between adipocytes and fibroblasts is important in contexts beyond wound healing. Fibrosis is a condition of excessive extracellular matrix deposition that is common in autoimmune disease, cardiovascular and lymphatic disease, and most forms of organ failure. The replacement of adipose tissue with fibrotic matrix is seen in many of these conditions, and it is a hallmark of the autoimmune disease scleroderma (Fleischmajer et al., 1971). Lineage tracing in a bleomycin model of skin fibrosis supported the idea that adipocyte dedifferentiation could be a source of pro-fibrotic fibroblasts or myofibroblasts (Marangoni et al., 2015). Further study of cells undergoing the adipocyte-to-fibroblast transition is warranted because of its potential impact on understanding common fibrosis mechanisms.

Adipocyte dedifferentiation is suggested to contribute to wound healing in two ways: first by releasing lipids that promote macrophage infiltration necessary for early wound healing, and second as a source of myofibroblasts that generate and remodel scar tissue (Shook et al., 2020). However, the process of adipocytes changing cellular identity into fibrogenic cells has been called into question (Kalgudde Gopal et al., 2023). Our results demonstrate that most *Pdgfra*^+^ adipocytes remain at the wound edge and do not migrate into the wound bed, but we also found αSMA expression in labeled cells at the wound center. Adipocytes are generally understood to be a highly plastic cell type (Roh et al., 2020; Rosenwald et al., 2013; Shen et al., 2022; Shook et al., 2020; Song and Kuang, 2019; Wang et al., 2018; Zhang et al., 2019; Zhu et al., 2022). Regardless of whether adipocytes can become true myofibroblasts, the number of adipocyte-lineage cells that enter the wound bed is tiny compared to the large number of fibroblasts and myofibroblasts originating directly from fibroblasts. Therefore, it does not seem possible that adipocytes significantly contribute to the cell population that builds and remodels scar tissue in typical skin wounds. If dedifferentiating adipocytes are not significant to the scar forming population, why do they proliferate? One possibility is they proliferate to serve as a latent source of adipogenic cells for reconstituting dermal fat after wound healing. Dermal fat regeneration does not occur in 5mm wounds that we employed, but it does occur in larger wounds in response to hair follicle regeneration, and a myofibroblast origin was implicated by lineage tracing (Guerrero-Juarez et al., 2019; Plikus et al., 2017). Learning whether dedifferentiated adipocytes can re-differentiate into adipocytes will require further studies applying the large-wound model to APαT mice.

Adipocyte dedifferentiation in wound healing requires the lipolysis enzyme adipose triglyceride lipase (Atgl, encoded by *Pnpla2*) (Shook et al., 2020), but otherwise how this process is regulated is unexplored. We sought to characterize PDGF and TGFβ as potential regulatory signals. This hypothesis was based on the general pro-fibrotic and anti-adipogenic properties of PDGF and TGFβ receptor signaling (Iwayama et al., 2015; Roh et al., 2020; Sun et al., 2017; Sun et al., 2020) as well as the upregulation of *Pdgfra, Pdgfrb, Tgfbr1, Tgfbr2*, and *Tgfbr3* in dedifferentiated adipocytes (Shook et al., 2020). Using APαT mice to delete *Pdgfra* from adipocytes, we found that only 10% of *Pdgfra*^+^ adipocytes converted into fibroblasts, while expression of constitutively active PDGFRα caused 90% conversion. Similar to *Pdgfra* knockout, deletion of *Tgfbr2* produced a low percentage of conversion. Interestingly, *Tgfbr2* knockout had no effect on the expression of Tomato in APαT mice, indicating that TGFβ is not required for *Pdgfra* expression in wound edge adipocytes. To ask whether the high rate of conversion driven by constitutively active PDGFRα requires TGFβ signaling, we generated APαT mice with constitutively active PDGFRα and deletion of *Tgfbr2*. But the double mutant wounds still had a very high rate of conversion, arguing that TGFβ signaling is not required for the effects of strong PDGFRα signaling. We conclude that the two pathways promote dedifferentiation through largely independent mechanisms. We still do not know what signal or stimulus in the wound edge microenvironment activates *Pdgfra* expression in adipocytes that are specified to dedifferentiate. The Notch pathway and Yap/Taz are potential candidates, as both have been shown to promote dedifferentiation when ectopically activated in mice (Bi et al., 2016; Shen et al., 2022).

In this study we have not tried to describe an overall wound healing phenotype after manipulation of PDGF and TGFβ signaling in adipocytes. We previously showed that fibroblast-specific knockout or gain-of-function of *Pdgfra* has a major effect on fibroblast proliferation and differentiation into myofibroblasts, which significantly impairs wound contraction and re-epithelialization (Yao et al., 2022). But the role of adipocytes in wound healing seems distinct from that of the local fibroblasts. Impairing lipolysis by adipocyte-specific *Pnpla2* deletion has transient effects on the inflammatory stage of wound healing without affecting fibroblast recruitment and re-epithelialization of the wound (Shook et al., 2020). We suspect that reduced adipocyte dedifferentiation because of loss of PDGF or TGFβ signaling could have similar effects as ablation of lipolysis via *Pnpla2* deletion.

In conventional single-step lineage tracing with tamoxifen-regulated CreER or doxycycline-regulated Cre, a reporter gene can usually be detected within 24 hours of drug treatment and sometimes earlier. The high specificity of our multi-step system has a limitation in that it takes time for the reporter to “develop”. Tamoxifen primes the reporter by allowing *Pdgfra* to express Flp°, but this does not occur until *Pdgfra* expression occurs. A major determinant of the reporter’s timing is how quickly after wounding the promoter can ramp up Flp° expression to a threshold that efficiently recombines R26-fSf-tdTomato. We used codon-optimized Flp° that is equivalent to Cre in its activity in mammalian cells (Raymond and Soriano, 2007). Nevertheless, we could first detect the reporter in APαT mice on day 4 after wounding. This means that the first 96-hours of adipocyte dedifferentiation could not be visualized by our system. Also, APαT mice are presumably most effective for tracing adipocytes that induce high *Pdgfra*^*Flp*^ expression, which helps explain why many wound edge adipocytes remained Tomato^neg^. As we showed previously, mosaicism is inherent to, and sometimes an advantage of, tracing with sequential activities of inducible Cre and Flp (Sun et al., 2020). We expect APαT mice to be useful for specifically tracing adipocyte dedifferentiation in other conditions, such as, hair cycling and skin fibrosis (Marangoni et al., 2015; Zhang et al., 2019), lactation (Wang et al., 2018) and tumor stroma development (Zhu et al., 2022).

## METHODS

### Animals

B6;129-Tg(Adipoq-cre/Esr1*)1Evdr/J mice (AdipoqCreER) (Strain #024671) and B6;129-*Tgfbr2*^*tm1Karl*^/J mice (Tgfbr2^floxed^) (Strain #012603) were purchased from The Jackson Laboratories. Pdgfra^Flp^ and Pdgfra^K.Flp^ mice were developed in the Olson laboratory and R26^Frt-STOP-Frt-tdTomato^ mice were derived from Strain #021875 as described previously (Sun 2020). All animal experiments were performed according to procedures approved by the Institutional Animal Care and Use Committee at the Oklahoma Medical Research Foundation. Mice were maintained on a 12 hr light/dark cycle and housed in groups of two to five with unlimited access to food and water. All strains were maintained on a mixed C57BL6/129 genetic background at room temperature. Both males and females were analyzed.

### Wounding

Tamoxifen was prepared as a 20mg/mL stock in corn oil and mice were gavaged three times on alternating days with 100 mg tamoxifen/kg body weight. Wounds were created four days after the last Tamoxifen treatment. To create excisional wounds, 7-9 weeks-old mice were wounded during the telogen phase of hair cycling. Mice were administered analgesic (Ketoprofen 5mg/kg) followed by inhaled anesthesia (5% isoflurane/1% oxygen). Dorsal hair was shaved and then completely removed using depilatory cream. The exposed skin was sterilized with 70% ethanol. Excisional wounding was performed using a 5mm biopsy punch to create 4 full-thickness dermal wounds. Mice were then single housed and wounds were left uncovered during healing. At the time of harvest (4-7 days post wounding), wound areas and a margin of unwounded skin were harvested and imaged from the underside with a fluorescent dissecting microscope. Wounds and fat depots were then fixed overnight in 10% neutral buffered formalin or for 1 hour in 4% paraformaldehyde. For *in vivo* proliferation assays, mice were intraperitoneally injected with 200 μL of 2 mM EdU solution in 0.9% saline 24 hours and 4 hours before sacrifice.

### Staining and imaging

Mouse wounds and fat depots were embedded in optimum cutting temperature compound and frozen sections were cut. Some wound beds were cut in a cross-section (or full-thickness) orientation at 14 μm thickness. Most wound beds were cut in a horizontal orientation, parallel to the plane of the skin at 20 μm thickness, because this allowed a larger area of the dermal adipose layer to be seen in a one section. Fat depots were sectioned at varying thicknesses depending on adipocyte size: visceral/perigonadal fat at 100 μm, subcutaneous/inguinal fat at 50 μm, dermal fat in a horizontal orientation at 20 μm, and brown/interscapular fat at 14 μm. Sections were stained with antibodies listed below, counterstained with DAPI, and mounted in Fluoro Gel with DABCO. Images were captured on a Nikon Eclipse 80i microscope or Nikon AXR confocal microscope. For EdU detection, sections were incubated with EdU reaction cocktail (175 μL PBS, 4 μL CuSO4, 0.2 μL Alexa Fluor 647 Azide, and 20 μL freshly prepared 0.5M ascorbic acid) for 30 minutes.

### Image quantification

Quantification of Plin1 and Tomato at the wound edge was performed on multiple horizontally-cut sections for each wound. Between 71 – 281 Tomato^+^ cells were scored at the wound edge for each mouse. At the wound center where labeled cells were relatively rare, between 14 – 29 Tomato^+^ cells were scored per mouse. For each mouse the % Plin1^+^ cells among the Tomato^+^ population, or the dedifferentiation index, was calculated by dividing the number of Plin1^neg^Tomato^+^ cells counted by the total number of Tomato^+^ cells counted. Quantification was performed manually. Some experiments were blinded, but in many cases, it was not effective because obvious differences in labelling revealed the genotype.

### Statistics

To determine significance between two groups, comparisons were made using Student’s t-test. Analyses across multiple groups were made using a one-way ANOVA in GraphPad Prism.

### Antibodies

αSMA: anti-alpha smooth muscle actin antibody, rabbit, Cell Signaling cat #19245

PPARγ: anti-PPARγ antibody, rabbit, Invitrogen cat #MA5-14889 (requires 1 hour fixation in 4% PFA)

Plin1: anti-Plin1 antibody, rabbit, Cell Signaling cat #9349

Secondary antibody: anti-rabbit IgG Cy5-conjugated, Jackson ImmunoResearch cat #711-175-152

## Acknowledgements

We would like to thank Drs. Sathish Srinivasan, William Berry, and James Tomasek for their helpful suggestions on this manuscript. We also thank the Microscopy Core and Mouse Phenotyping Core Facilities (associated with P30-GM114731 and P20-GM103636) of the Oklahoma Medical Research Foundation Centers of Biomedical Research Excellence. L.E.O. is funded by NIH-NIAMS R01s AR073828 and AR080896; the Oklahoma Center for Adult Stem Cell Research – a program of TSET; and the Oklahoma City-based Presbyterian Health Foundation.

## Author Contributions

**Longbiao Yao:** Methodology, Investigation, Visualization, and Writing – Review and Editing; **Sunhye Jeong:** Investigation, Validation, Visualization, Validation, and Writing –Review and Editing; **Hae Ryong Kwon:** Visualization, and Writing – Review and Editing; **Lorin E. Olson:** Conceptualization, Methodology, Investigation, Visualization, Funding acquisition, Supervision, and Writing – Original Draft, Review and Editing.

---

**Supplemental Figure 1.**
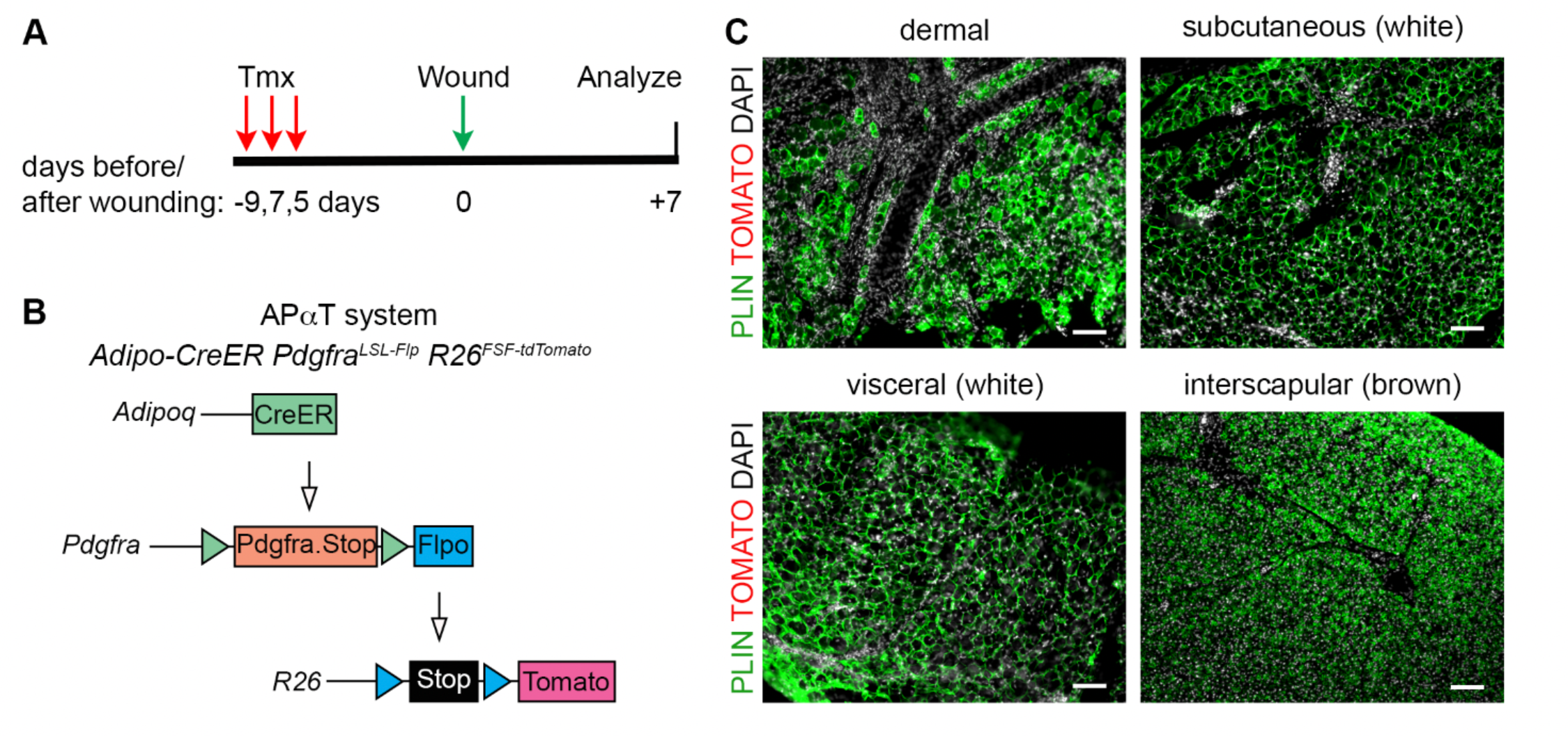
*AdipoqCreER* does not overlap with *Pdgfra* expression in unwounded fat. (A) Timeline for analysis of fat depots distant from skin wounds, which is the same timeline as Figure 1B. (B) Schematic of the genetic strategy for APαT, which is the same mice as Figure 1A. (C) Plin1/Tomato co-labeling of unwounded fat from APαT mice. There are no Tomato-labeled cells in unwounded fat. Scale bar 100 μm.

